# The role of cultural norms in shaping attitudes towards amphibians in Cape Town, South Africa

**DOI:** 10.1101/681403

**Authors:** Peta Brom, Pippin Anderson, Alan Channing, Leslie G. Underhill

## Abstract

Urban ecosystems are increasingly viewed as an important component within strategies for wildlife conservation but are shaped as much by natural systems as they are by social and political processes. At the garden scale, attitudes and preferences govern design and maintenance choices including the decision to encourage or discourage specific faunal presence. At the global scale, charismatic taxa that are well-liked attract more conservation funding and volunteer stewardship. Amphibians are a class of animals that are both loved and loathed making them a suitable subject for comparing and unpacking the drivers of preference and attitudes towards animals. We conducted a mixed methods survey of 192 participants in three adjacent neighbourhoods in Cape Town, South Africa. The survey included both quantitative and qualitative questions which were analysed thematically and used to explain the quantitative results. The results revealed that attitudes formed during childhood tended to be retained into adulthood, were shaped by cultural norms, childhood experiences and the attitudes of primary care-givers. The findings are significant for environmental education programmes aimed at building connectedness to nature and biophilic values.

## Introduction

With more than half the world’s human population urbanized [1], urban environments are the only place where many people will have opportunities to experience nature and urban nature is fundamentally shaped by the choices people make. Levels of biophilia, attitudes and perceptions as well as perceptions of nature result in certain species being prioritised for conservation, whilst less charismatic or “liked” species attract smaller budgets and less research attention [2]. Additionally, social norms [3], individual preferences [4], attitudes [5], perceptions [6], cultural beliefs [7], and even identity [8,9] can result in different landscaping practices and a desire to cultivate and attract or remove and deter one species over another. What we like and don’t like therefore matters to the future of urban nature conservation.

Amphibians have ecological importance in many ecosystems around the world. They are an essential link in the natural food web and are important bio-indicators in determining wetland and river health whilst regulating invertebrate populations [10]. They are also the most threatened vertebrates on earth with approximately 41% of the entire class recognised as such [11]. The most widely attributed reason for amphibian decline is habitat loss associated with land use changes and development [12], but none of the factors associated with agriculture and urbanisation can readily account for the declines that have been found in areas apparently unaffected or remote from land-use change. Declines occurring in remote areas are instead attributed to climate change, UV radiation, and diseases such as ranavirus and chytrid fungus [13]. The spread of these diseases is facilitated by species invasions and climate change [13]. Climate change is an emerging land-cover change driver. Predictions indicate shifts in natural habitats 80 years hence will occur at rates 500 times faster than current trends [14]. In short, amphibian species are threatened globally by a changing world and these changes are anthropogenic.

Cities are arguably the most altered sites of change. Urban environments are prone to urban warming, and local climate changes within cities have occurred at faster rates than surrounding areas [15] yet Ives *et al.* (2015) found that Australian cities consistently supported a greater number of threatened species than “all other non-urban areas on a unit-area basis” [16] and Goddard *et al.* (2010:90) recognise that “globally declining taxa can attain high densities in urban habitats” indicating the need for a reassessment of the value that urban ecosystems can contribute towards conservation. For amphibian populations, a large-scale citizen science study in North America found that although urban populations of amphibians are smaller than their wild counter-parts, they appear to be declining at similar or slower rates [17], suggesting that urban environments may be able to provide refuge for some species of amphibians if conditions are suitable.

In many cities around the world, retention ponds, attenuation ponds and rain-gardens, developed as components of stormwater systems, have been colonized by amphibians as breeding habitat [2,18–20]. Studies which focused on urban ponds have found that natural urban wetlands and constructed habitats have similar occupation [21], but that the quality of terrestrial habitats is as important to amphibians as the in-pond conditions [22,23], highlighting the fact that amphibians rely on both aquatic and terrestrial habitats.

Amphibians are both loved and loathed, making them a suitable class for unpacking human attitudes. All over the world they are steeped in myths and superstitions that have been brought to us through time. Walter Rose (1929) attributed the mythologies he encountered several typical characteristics of the class. The metamorphosis process where tadpoles visibly grow legs before leaving the pond associates them with transformation. Frogs are seemingly magical in their ability to crawl into tiny cracks and burrows. They disappear for months during aestivation and then, during a storm they can seem to appear from nowhere – leading to myths about frogs raining from the sky [24]. In Western society, frogs are associated with magic and metamorphosis such as in the image of the Frog Prince as documented by the Grimm’s brothers and popularized by Dysney, and that of Shakespeare’s famous witches brew (Macbeth), which included “Eye of newt and toe of frog”. Tarrant *et al.* (2016) explain some of the mythology as an inability for many people to make sense of amphibians as animals. This is reflected in stories where frogs are turned into human-like creatures with mystical powers.

South Africa is a diverse country of many cultures and heritages and 11 official languages. Some groups hold beliefs which fuel negative response attitudes towards amphibians. For example, amphibians were documented as the second most feared animal amongst 120 Zulu respondents across various age groups (snakes were the first) and this fear often led to direct killing of amphibians [2]. There are four major ethnic divisions among black South Africans, namely the Nguni, Sotho, Shongaan-Tsonga and Venda. The Nguni is the largest and can be divided further into four groups, of which isiZulu is Northern and Central Nguni, isiXhosa is spoken by Southern Nguni, Swazi by those from Swaziland and Ndebele in the Northern Provence and Mpumalanga. isiXhosa and isiZulu are particularly similar and share similar customs. The clear-cut distinction made today between isiXhosa and isiZulu originated in colonial distinctions between Natal and the Cape and later intermarrying and cultural borrowing from the Khoikhoi amongst Southern Nguni cemented these distinctions [25]. These groups share a rich oral tradition as a primary means of memory retention and heritage [26] so beliefs about animals are often passed down between generations.

Tarrant *et al.* (2016:1) note, “That the average amphibian receives 75% less funding than the average listed mammal, bird or reptile, and 90% less funding than the average listed fish reflecting the less-popular status of amphibians in general” [2]. One of the effects of ubiquitous negative attitudes is that it translates into lower prioritisation for conservation. It therefore becomes important to focus on the ways that attitudes are shaped and influenced if conservation efforts are to gain the traction required from the public to reach their targets. This study explored the themes driving attitudes to amphibians in a neighbourhood composed of three adjacent suburbs in Cape Town, South Africa.

## Methods

A survey by questionnaire (See addendum A) was developed in order to cover four areas likely to correlate to attitudes, namely, demographics, preferences, knowledge and personal childhood experiences. The initial question set was adapted from Tarrant *et al.* (2016) who aimed to test knowledge, beliefs and liking amphibians. Tarrant *et al.*’s (2016) questions were all measured on a 10 point Likert scale, whereas we asked instead that respondents select from a list which best describes the feelings towards amphibians with choices between, ‘I like frogs’, ‘frogs are ok’, ‘frogs are gross’, ‘frogs are scary’ and ‘I have no feelings about frogs’. Although blunt, this self-identified response held valid as a position and framework throughout the cases. Cultural belief questions were used in the same format as Tarrant *et al.* (2016), whereas knowledge questions were drawn from both Tarrant *et al.* (2016) and added to from du Preez *et al.*’s (2009) introductory section on amphibians resulting in “Frogs/toads are considered harmless to people” and “Some frogs/toads secrete a mild toxin on their backs as a defence mechanism (e.g. when hurt).” [27]. Preferences questions were added based on the work of Belaire *et al.* (2015), who measured residential preferences towards birds. This produced questions that asked respondents to agree or disagree on a five-point scale with the statements “I like listening to frog/toad calls when it rains” and “Frog/ toad calls keep you awake at night.”

In order to relate the questions to de Groot *et al.*’s (2003) preference ladder [29], questions were designed to consider behavioural responses at scales within the home by asking respondents first what respondents would do if they found a frog in their garden and then if they found it in their homes. Respondents were also asked if they thought amphibians should be protected in the wild and then in green spaces in the city. To test the specific levels of preference, respondents were asked to look at four images of amphibians that each represented the typology of a. rain frog, b. reed frog, c. toad and d. river frog to determine how attitudes to specific types of frogs would differ from general ideas. The frogs selected for the images are native to the City of Cape Town and could be encountered in resident’s gardens (Figure 1)

**Figure 1.**
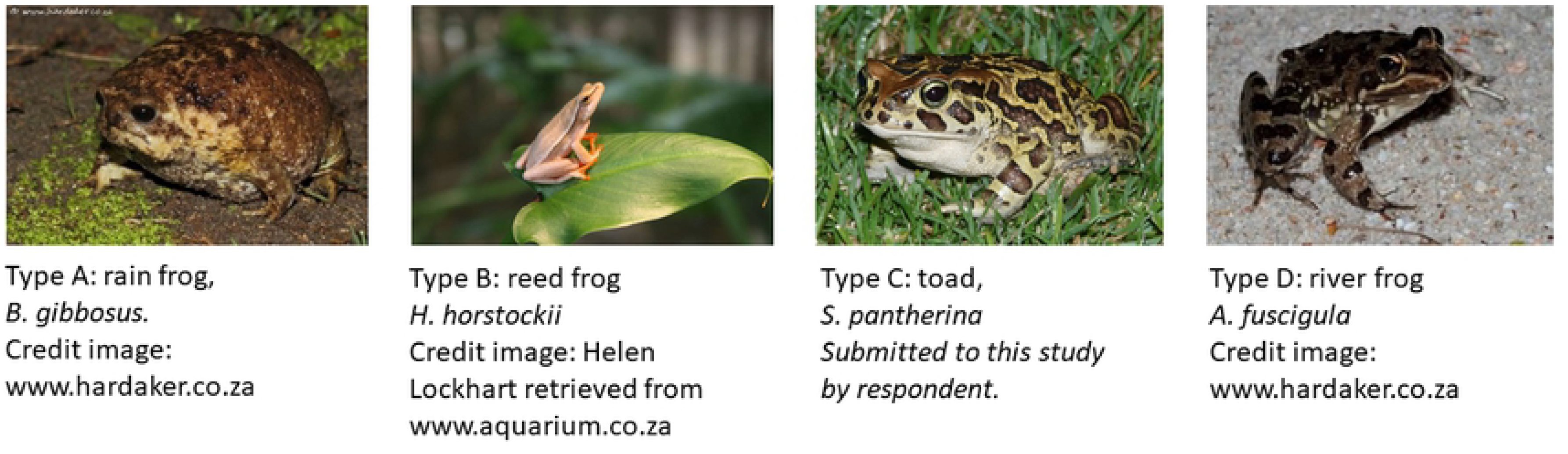
Flash-cards used to measure specific attitudes towards different frog types as would occur within the City of Cape Town

Tarrant *et al.* (2016) speculated that those who had positive experiences of frogs in their childhood at an age younger than 10 were more likely to have a strong affiliation towards frogs and so in order to explore the relationship between childhood experiences, cultural beliefs and attitude towards frogs, respondents were asked “Do you have any strong memories of coming into contact with frogs from your childhood, or any memories of something that someone, a parent or teacher, told you about frogs that you would like to share?”

Prior to commencing field work, ethical clearance was sought and granted by the University of Cape Town under approval code FSREC 021 - 2016.

36 respondents were visited, and questionnaires were administered directly. Where possible both gardeners and home-owners were interviewed. During this time the questions were fine-tuned both in the phrasing and prompting. The questionnaire was then converted to a digital format using survey monkey and a link was posted to social media groups. To ensure the inclusion of those without access to digital platforms, administration was undertaken on the street after the digital platform was made available, offering respondents a choice of platform for engagement. Posters and flyers were printed inviting respondents to find the survey questionnaire online. The posters were put up in local restaurants, bars and teahouses in Rosebank, Little Mowbray and Observatory.

The team was assembled from Environmental and Geographical Sciences undergraduates at UCT and comprised of five women, three of whom were isiXhosa first-language, one was Kenyan and one was South African English first-language. Four members of the team stood for one week-day morning on Mowbray, Rosebank and Observatory railway station platforms between 7:45 and 9:30am and interviewed commuters leaving the respective suburbs. Flyers were also handed out inviting commuters to logon using their phones during their train-ride. On a Sunday morning two members of the team went to the village green in the centre of Observatory and interviewed 15 street-dwellers who had come to take advantage of a soup kitchen that would be setting up later in the day. A team of three visited the Observatory Library on a Wednesday morning.

Recognising that street harassment and begging are problems in these areas, the team wore matching T-shirts with bold print that said “Urban Biodiversity Research” on the back, thereby announcing the team’s intention and legitimacy. Overall, the community was receptive, and we were received with a mixture of enthusiasm, curiosity and tolerance.

## Analysis

Results were processed descriptively (counts, percentages, means and standard deviations) then cross-tabulated to explore the relationships between demographics and attitudes and preferences, then knowledge and beliefs, responses to amphibian presence in the garden and home and finally the relationship between childhood memories and disposition towards amphibians was assessed. Associations were evaluated using Chi-squared test and one-way ANOVA between disposition towards amphibians and demographics, attitudes, preferences knowledge and beliefs. Correspondence analysis was performed to explore the relationship between disposition towards amphibians and themed narratives visually.

## Results

A total of 192 survey responses were obtained. The respondents were predominantly between the ages of 18 and 50, with less than 5% falling below 18 years of age and above 70. The majority of respondents (57%) said that English was their mother tongue reflecting the dominant demographic of the area. isiXhosa (19%) was the second language group in the respondent set, whilst the remainder self-identified as Afrikaans (3%), Bilingual Afrikaans-English (3%), isiZulu (2%), and Other (15%) which included a group of nine international languages from African and European countries of origin.

Overall education levels were high. 60% (n=113) of respondents had completed at least some form of higher education, reflecting both the dominant age-groups of the interviewees and the education levels of the suburb due to its socio-economic status and proximity to tertiary educational institutions. It may also reflect a response bias of willingness to engage with research from those with higher education. 69% (n=129) of respondents liked frogs or said they were ‘OK’ whilst 10% (n=18) had a neutral response and 21% (n=40) had a negative response, saying they were ‘scary’ or ‘gross’.

Those that liked frogs tended to leave them alone, remove them from their houses to the garden or to a lesser extent take them to the river or nearest wetland and release them. In these instances, the reason given for removal from property was due to perceived threat from pets, or the perception that the frogs were not in their natural or preferred habitat. Those that did not like frogs would either call someone to remove them from the property or respond in fear.

The majority (89.5%, n=162) agreed that frogs should be protected in the wild but protecting them in the city came into competition with other objectives including access for leisure and social pursuits. In this instance, respondents asked if protecting them would compromise their ability to use green spaces freely and asked for clarity on what was meant by “green spaces” expressing uncertainty, the definition given covered public open space and corridors. In spite of the hesitation in the tone of the interviews, 83% (n=161) of respondents agreed that frogs should be protected in green areas within the urban edge. Respondents were more ambiguous about making it easier for frogs to move through the city, citing feasibility as the main concern and prioritized human needs within the urban and city space. When prompted with the statement that there may be simple cheap ways to improve mobility, 65.6% (n=118) of respondents agreed or strongly agreed that we should make it easier for frogs to move through the city. Those that did not like frogs tended to express the view that frogs should stay in the “wild” and disagreed with this statement.

### Language and Culture

Language was used as a proxy for culture and so is discussed in terms of culture. Of the isiXhosa-speaking respondents who said they disliked frogs, a cultural belief was reported that individual frogs found on their property out of the rainy season were sent by witchcraft to curse you. The remedy is to kill the frog, preferably by sprinkling salt on its back and then sweeping up the body. These qualitative responses were revealed in the coding of the “other” answers to the question “if you found a frog in your house, what would you do?” isiXhosa-speakers were most likely to report being phobic of frogs to the extent that they were unable to look at the flash-cards of examples of frogs. A few respondents reported a shift in attitude with urbanisation or gaining education.

**Figure 2.**
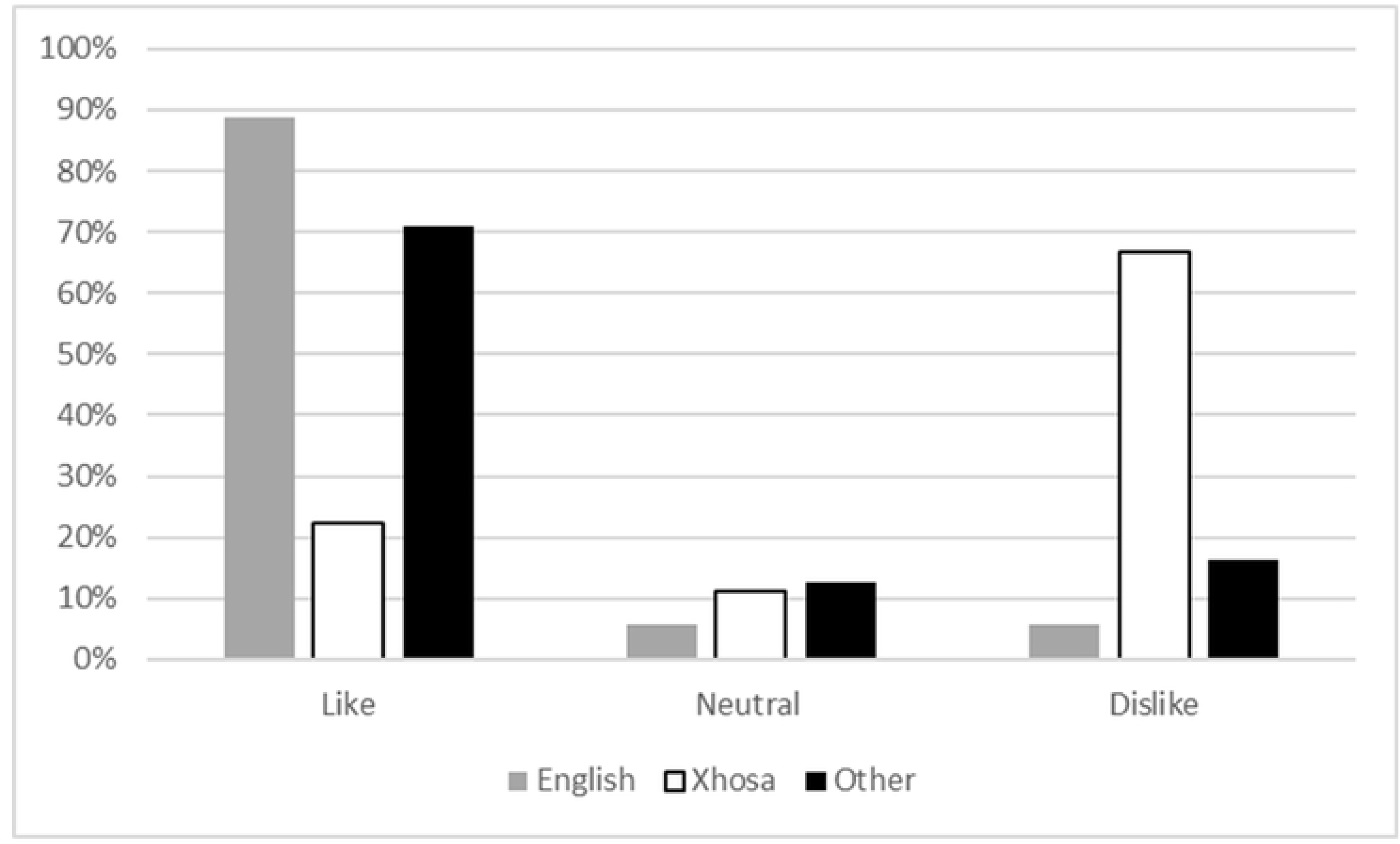
Feelings towards frogs split by dominant language groups

### Knowledge and Beliefs

The knowledge and belief scores were cross-tabulated attitudes (Table 3.1). The knowledge of those who liked frogs was significantly better (more accurate) than the knowledge of those who disliked frogs (two sample t-test t=5.99, d.f.=161, P<0.001). This is also reflected in the knowledge means of the three groups. Knowledge means were lower in the group that were afraid of frogs. Therefore, a correlation between positive attitude towards frogs and higher knowledge scores demonstrates that those that like frogs have more accurate knowledge of them. It does not however appear to be a causal relationship as people who like frogs may be inclined to search out accurate knowledge about them as much as those that have more accurate knowledge about frogs may develop an interest and affinity towards them. When cross-tabulated against education achieved, those with post-graduate education had lower knowledge and belief scores than those with undergraduate education, but both groups had higher scores than the group with secondary education. This drop in post-graduate scores can be attributed to the fact that those with research training tended to refuse to guess their answers choosing rather to say they did not know. Table 1 presents the median scores for each preference group.

**Table 1.** Cross-tabulation of means for knowledge and belief scores against attitudes towards amphibians.

| Feelings<br>(collapsed<br>categories) | Knowledge<br>Cat. Score | Beliefs<br>Cat. Score | Total<br>Knowledge<br>Scores |
| --- | --- | --- | --- |
| 1 Like | 2.93 | 2.56 | 5.49 |
| 2 Neutral | 2.18 | 1.94 | 4.12 |
| 3 Dislike | 2.21 | 1.66 | 3.87 |
| Total (average) | 2.71 | 2.31 | 5.02 |
| N | 180 | 180 | 180 |
| Std. Deviation | 1.05 | 0.886 | 1.636 |

### Specific Preferences

The most popular frog was *H. horstockii* which was reported by 76.64% of respondents as being ‘likeable’. This was followed by the *A. fuscigula* (55.3%), the *S. pantherina* (54.8%) and finally, *B. gibbosus* at (32.4%). The results are presented figure 3 and show specific attitudes towards individual species differs from the general conception of “frogs” as an animal.

**Figure 3.**
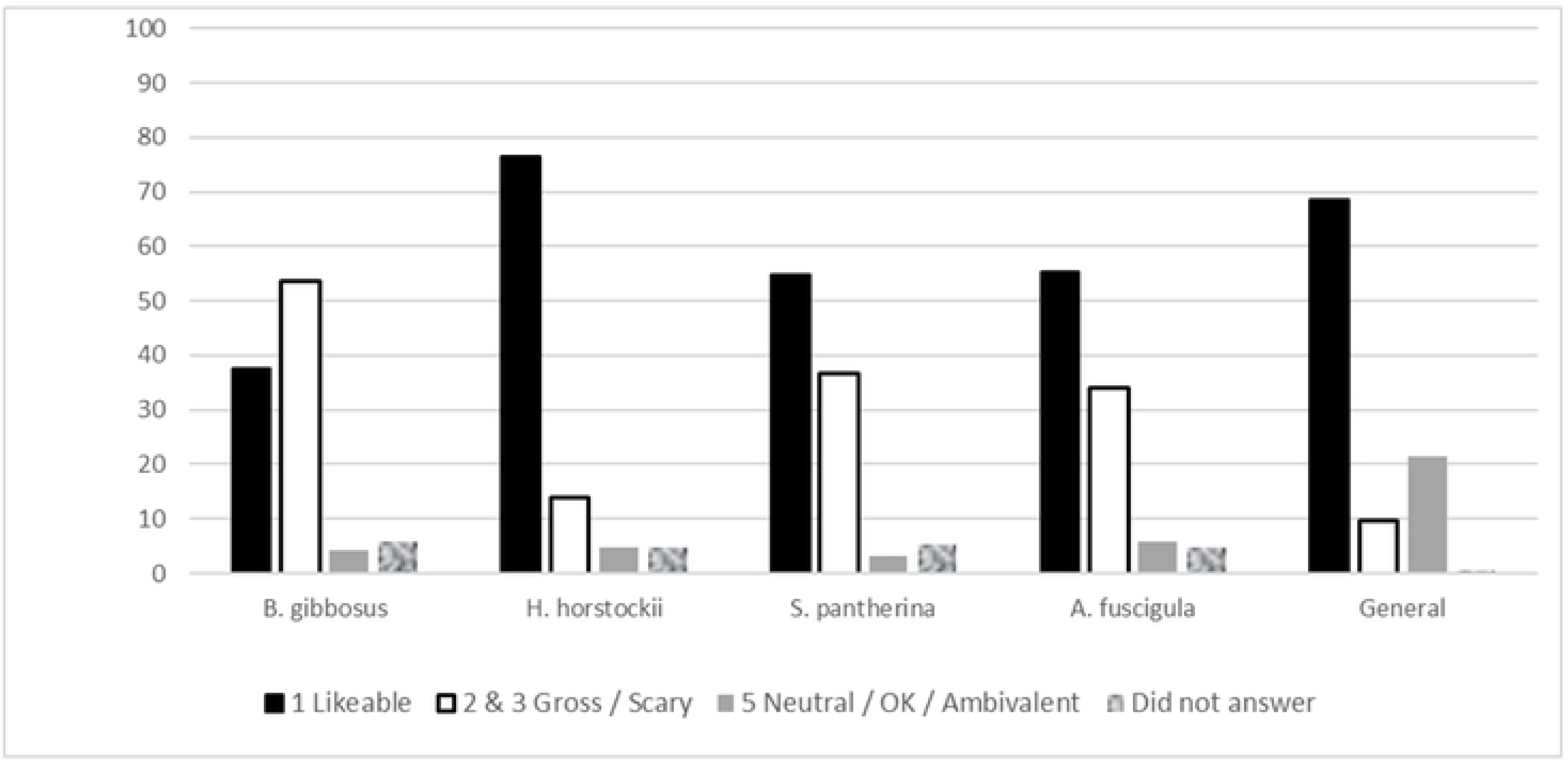
Frequencies of preferences towards different frog species B. gibbosus, H. horstockii, S. pantherina and A. fuscigula compared to general attitudes towards frogs.

### Behaviour at Spatial Scales

Behaviour responses did not change significantly between the house and the garden. The exception was for those who said they would try to find out more about the frog if it was in the house (19.4%) and those who said they would leave the frog alone if it was found in the garden (55%). 60% of all respondents said that they would remove frogs from the house and put them in the garden (or call someone to do so); whilst 12.9% of the sample said that they would kill it, put salt on it or chase it away. Only 4.3% said they would leave it if it was found in the house. 12.4% of the sample would remove the frog from the property or take it to the river, either due to the perception that the river was where it belonged, out of concern for feline predation or due to fear and disgust.

### Life Experiences

Responses to the question “How old were you the first time you remember coming into contact with a frog?” (figure 4), fell into the following categories; i. did not know or couldn’t remember (n= 21); ii. under the age of five (n= 93) or iii. between the age of six and 10 (n=61). Only a few outliers within the sample did not have recollection of some contact with frogs before they were ten years old (n=13). When the age of recollection of first contact with frogs was cross-tabulated with attitude towards frogs, the proportion of those that dislike frogs peaked in the 6–10 age category, and the proportion of those that liked frogs peaked in the 0–5 age group. Having said this, the samples have large overlapping areas indicating both positive and negative outlooks within both age groups.

**Figure 4.**
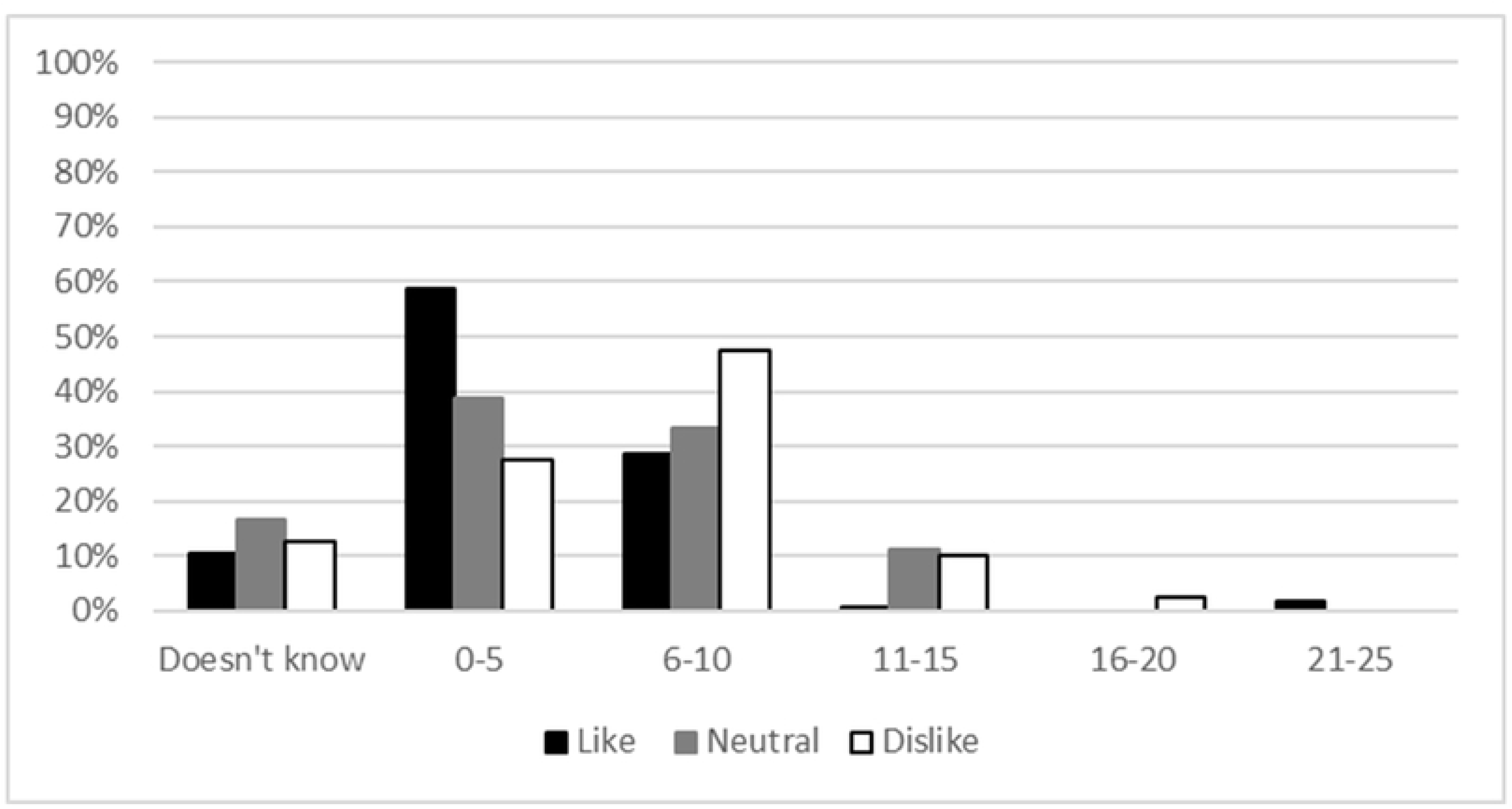
Respondent recollection of age when they first saw or came into contact with a frog.

The thematic analysis of the narrative of a memory from childhood show clear distinctions between those that find frogs ‘gross’ or ‘scary’ and those that find them ‘likeable’ or ‘OK’. Those that have no feelings did not reveal any clear consistency in themes, but 61% (n=11) of them had no recollection to share. Catching tadpoles (n=19) featured frequently as a theme amongst those that had an affinity for frogs. The second theme was childhood discovery (n=14) recounting playing with and discovering frogs, seeing them or hearing them, often characterised by a sense of wonder. Respondent #129, who liked frogs said “*I remember at [place name] as a child I went to the garden, playing with water and a lot of tiny frogs popped out and I was so amazed and held them on my hands.”* Respondent #93 said *“looking for frogs on the sides of mountain pools (often around* Disa uniflora*) after a long hot walk on the mountain. If you could stay in the cold brown water long enough, we used to see how close we could swim to them before they jumped into the pool.”*

Parental biophilia also featured among this group (n=5), in which a primary care-giver would tell the child not to harm the animal or would be involved in facilitating the interaction, either by instruction or taking them frogging. Respondent #4 said *“My dad calling us all into the garden at night to show us a leopard toad by torchlight. It happened fairly often! And then I did not see one for years until about 12 years ago in our [place name] Garden …a long space in between!”*, Some (n=5) reported trying to keep them as pets, and some reported playing with them more destructively, or using them to play practical jokes on their friends (n=7) *“I once found a frog and put it in my sister’s room and she freaked out.”*, others remember listening to them during the rain or at night (n=4), and lastly, there were those who witnessed the killing of frogs with some distress, implying that they were already familiar with them, were unafraid and held some empathy (n=5). These themes and accounts had in common direct interaction, fond recollection and that the adults either facilitated, or allowed engagement with minimal interference or warning.

On the other hand, those that reported fear of frogs tended to hold beliefs in the ability of amphibians to harm them. Two main themes emerge. Firstly that they were told by an adult or parent, that touching them (or even looking at them in one case) can result in severe rashes or infections (n=7) and secondly that frogs are associated with witchcraft (n=6), Respondent #188 said *“Where I come from, some people say frogs are sent by witchcraft, especially if it’s not raining or it’s unseasonably dry.”* Additionally, those that had been chased with frogs or startled also featured (n=6). Respondent #83 who thought frogs were ‘scary’ said "*Someone put it on me and I ran away and that’s when I knew I was scared”*.

Respondents across the like-dislike spectrum described frogs coming into the house, out of the ground, out of the drains in large numbers. One respondent who liked frogs said *“I remember living on an old farm in [place name] and one very rainy, stormy night we woke up to hundreds of frogs popping up from under the floorboards and trying to put on a pair of my mom’s high heels to avoid them jumping on my feet.”*

Figure 5 presents the correspondence analysis between the narrative themes and the attitude and illustrates the clustering of narratives that documented experiences, role-model attitudes and cultural beliefs with categories of attitudes and feelings towards amphibians in general. The model is statistically significant with the chi-square value at 86.295 (df=36) and p= 0.000. Dimension 1 shows the correspondence between the attitudes ‘dislike’, ‘neutral’ and ‘like’ and the themes found in the narrative. The theme ‘startled’ is an outlier on dimension 2, because respondents with a memory of being startled by a frog had varying attitudes depending on the context of the story and factors recorded in the other categories such as cultural background and parental biophilia. The close clustering of the ‘like’ and ‘dislike’ themes on both dimensions indicates the strength of the correspondence between the memories and the attitude.

**Figure 5.**
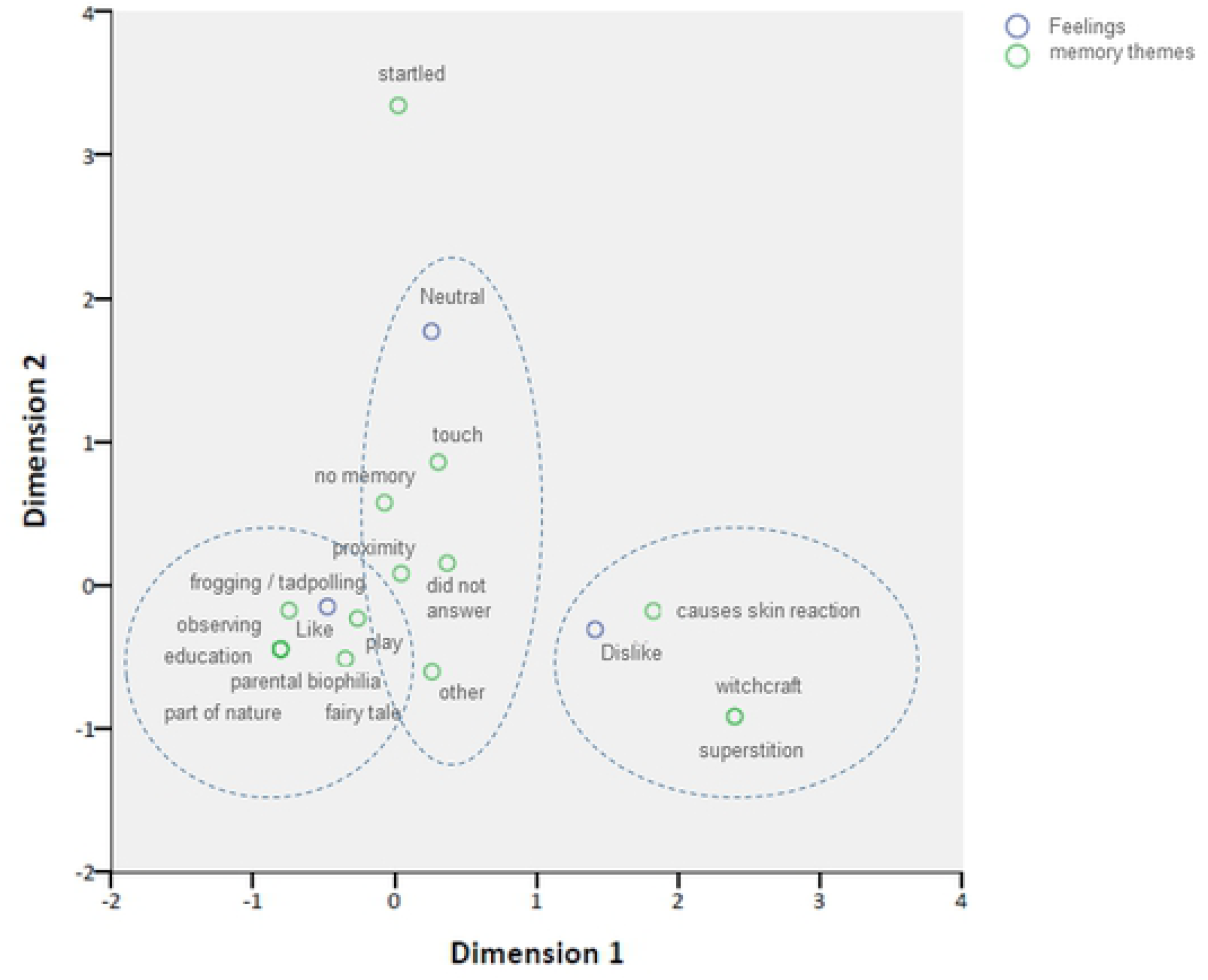
Correspondence analysis of narrative themes and attitude towards amphibians.

## Discussion

This study examined the preferences of a Cape Town community towards amphibians and explored attitudes using a composite approach drawing from a number of sources. De Groot *et al.’s* (2003) preference ladder was used as a theoretical framework for exploring preferences. The findings of this study were consistent with those of de Groot *et al.’s* (2003) in that general preferences at the broad conceptual level can be different to the specific level [29]. This study compared general preferences to specific preferences in terms of space, behaviour and individual species. When asking about individual species, the arum reed frog, *H. horstockii* was much more popular than other species and many people who were generally afraid of frogs said they thought it was ‘likeable’ and were more likely to leave it alone if they found it in their garden. Knight (2008) found that people preferred animals that were ‘cute’ with more human-like proportions to their faces and proportionately larger eyes. The arum lily frog is smaller than the other species presented and has a smoother pattern (as opposed to the mottles, warts and striking patterns of the other 3 species) and softer colouring to it (white, cream and beige as opposed to dark browns and kakis). It was described specifically and variously as being ‘beautiful’, ‘elegant’, ‘harmless’ and ‘it looks poisonous’. In contrast, those with strong dislike or fear of the other species often compared the appearance of the skin to a snake. Those that liked the rain frog, tended to laugh at it and see it as ‘funny’ or ‘grumpy’, personifying its ugliness into something relatable. As the least popular frog, the rain frog’s image was often met with dismay and exclamations of “What is that?!” and “is that even a frog?!” The results of this study suggest that reasons for liking an individual species correlate with aesthetic appreciation and relatability. This is consistent with Knight’s (2008) findings that personification and relatability feature highly in the likelihood that individuals will respond to calls to champion a specific creature for conservation [30]. The findings suggest that it is easier to promote urban biodiversity using charismatic species such as has been argued by Goddard *et al.* (2010) and Knight (2008), however the differentiation between the specific and the general means that it may only improve attitudes towards an individual species without necessarily affecting overall attitudes to amphibians in general [30,31].

Impacting the general preference level is more complex given the multiple social influences and individual life-experiences that shape human preferences towards nature. De Groot’s (2004) research closely associated a general preference for nature with a biophilic self-identity. Biophilia has a number of related concepts that closely align with an affiliation with nature [32] and underpin the framework of *Connectedness To Nature* (CTN) [33]. Although CTN was not directly measured by this study, the themes that emerged within the results are consistent with the themes underpinning CTN theory [34] and thus this framework is used for discussing the results of the general preferences towards frogs.

Positive conservation efforts within the urban context would require a shift towards a culture of pro-environmental behaviour. A predictive relationship has been demonstrated between biospheric values and pro-environmental behaviour [32]. Biospheric values are held when “People judge phenomena on the basis of cost or benefits to ecosystems or the biosphere” (Stern & Dietz 1994:70) and are a result of CTN [32]. CTN is a framework which measures an individual’s ability to see themselves as part of nature [32]. To harm a part of nature becomes synonymous harming oneself. Individuals who hold biospheric values are more likely to engage in pro-environmental behaviour [33]. Klassen (2010:10) summarised the interrelationships of concepts and precursors of CTN in terms of four underpinning pillars, namely, *lived experiences; encounters and conversations with passionate, caring or dedicated role models; cultural background; and prior knowledge*. This study has rendered similar findings in terms of the themes emerging from the results correlated with liking or disliking frogs in general and will be discussed below.

### Lived Experience

Early exposure to frogs was not, on its own, a key predictor in liking frogs as an adult because children who play in nature tend to encounter them. Instead, the quality of the interactions (often coupled with the attitudes of role-models facilitating those experiences) influenced attitudes. This builds on the concept of relational values as suggested by Chan *et al.* (2016). For example, Palmer *et al.* (1999) found that 75% of Canadians, and 71% of Australians selected childhood experiences in nature as the number one reason for personal responsibility being felt towards the natural world. Hunter and Brehm (2004) suggest that particular events during a youth’s life could result in environmental values being enhanced or altered depending on the values as being positive or negative and Lekies and Beery (2013) determined that children who collected natural objects as a child scored higher than non-collectors on a measure of connection to nature, which is corroborated by this research that found that tadpole collecting was a prominent feature amongst the stories that were related by the group that reported liking frogs [36–39]. However, it is perhaps not the collecting itself, but the time and quality of the experiences within nature required for collecting that builds the CTN. Wells and Lekies (2006) conducted an earlier study in North America that suggested that children who participated in both “wild” and “domesticated” nature were put on a trajectory towards environmentalism [40] and Klassen (2010) had similar results when he compared the experiences of rural children and urban children. Duerden and Witt (2010) found that children who engaged in direct educational experience were more likely to engage in pro-environmental behaviour after the course had ended [41]. Klassen (2010) also pointed to multiple or regular positive experience in nature [34]. The current research has produced some additional results that indicate that the attitude of the carer, or adult facilitating these activities, has a prominent role to play in this trajectory. Individuals that were actively discouraged from playing with, observing or going near to amphibians in early childhood, retained their fear into adulthood.

### Role Models and Parental Figures

The role of parent was often mentioned in the narrative results as someone who either passed on an attitude of affiliation for nature, a superstitious outlook or a set of warnings. Klassen’s (2010) summary of CTN theory included encounters with passionate role models including friends, family, teachers, community members, social movement leaders and writers. These role models can shape the kind of experiences and learning about nature that takes place through facilitated nature engagement (e.g. taking the family to the beach or leading a hike) or knowledge dissemination in all its formal and informal forms. Likewise Duerden & Witt (2010) explored the affective behavioural and knowledge retention of environmental impacts that were indirect (classroom based), direct (nature based) or vicarious (stories, plays and entertainment) [34,41]. Three different types of role-modelling can be identified, that of family and friends (home), that of teachers, educators and community leaders (community), and that of public figures (public). This research has highlighted the role of home-based figures in early childhood foundation years and noted that positive experiences tended to be imprinted at preschool age, whilst negative attitudes were associated with recollection from the primary school age. Having said that, Klassen (2010) emphasised that CTN was influenced by multiple positive lived experiences with passionate, caring role-models. A primary care-giver is a role-model who will be present on a continuous and regular basis. When children are encouraged and facilitated by adults to explore, play and engage with nature it enables sense of wonder and connection – a desirable precondition for establishing connection to nature [34]. This research recognises the importance of parental attitude in the formation and transfer of values and attitudes and suggests that further research is required to understand how to effectively shift whole-family attitudes by engaging both children and parents in positive nature experiences.

### Cultural Background

Cultural background includes cultural beliefs, values, attitudes and opinions of family and community members [34]. It is reinforced by the norms that are enforced by community members (injunctive norms) as well as what individuals observe or believe of others (descriptive norms). These find expression in community practices and role-model enforcement [3]. In this study, language was used as a proxy for cultural identity and showed stark differences between groups. One isiXhosa-speaking male even refused to participate in the study saying “Why do you want to know that? Everybody hates frogs” thereby revealing the descriptive norm within his group. isiXhosa people tend to hold the belief that frogs are dangerous and can spit a poison that causes infection in humans, therefore one should not touch them and should rather run away if you see them. Frogs are widely regarded by experts as harmless, however many frogs carry a toxin which they secrete when they are critically harmed. The banded rubber frog, is common in the Eastern Cape and Kwa-Zulu Natal, the regional areas where isiXhosa and isiZulu are most widely spoken respectively. It secretes a particularly irritating toxin which can result in rashes or vomiting if handled extensively by sensitive individuals [27]. The presence of this frog may go some way to explaining the belief that frogs can cause a rash through spitting. This belief seems to preclude children from early encounters with frogs and discourages them from playing too close to them, so they are unlikely to have positive life experiences with frogs and the resulting phobia, or disaffiliation, is carried through into adulthood.

Nassauer *et al*. (2009) suggested that recruiting community leaders or celebrities to champion pro-environmental behaviour can assist in fostering positive norms within a given society [3]. It is important that environmentalists are sensitive to the cultural beliefs and systems of the people that co-exist with the ecosystems they seek to conserve. Understanding the underlying suspicions, beliefs and impacts is an important step towards garnering support for conservation efforts. Further research may look at groups with a particularly negative outlook on an animal class, e.g. snakes, or frogs, and explore the qualitative themes amongst the minority sub-group who do like them. Put more specifically: what is different about the life experiences of those that like frogs within the isiXhosa group?

### Knowledge

The knowledge results within this study showed a correlation between accurate knowledge and beliefs and liking frogs. The group that disliked frogs had a lower mean score for knowledge and beliefs. It is not clear if lack of accurate knowledge was driven by disliking frogs or if disliking frogs meant that individuals were disinterested in accurate knowledge. Furthermore, those who reported direct positive experiences with frogs in their childhood also scored higher on knowledge and beliefs and may be a precursor to retaining accurate knowledge.

We did not seek to measure the impacts of educational strategies but rather to determine what the factors associated with a general attitude of liking frogs was. We confirmed that there is a relationship between knowledge and liking frogs. We observed examples of both intergenerational knowledge and the use of knowledge in better environmental decision-making. In the first instance the knowledge of others (role-models) is a factor in driving the value-basis during the formation/ deepening of CTN during childhood, whilst in the second instance, knowledge becomes a factor that shapes decision-making and pro-environmental actions. Therefore learning, whether formal, informal, direct or indirect is an integral foundation to fostering environmental behaviour, however it is not an actor that drives the formation of positive attitudes on its own. Thus it is important that quality information continue to be made regularly available to the public in order to facilitate appropriate pro-environmental behaviour and continue the cycle of generating experiences that drive biospheric values. Figure 6 below attempts to map out the relationships between the different aspects which work together to shape general attitudes.

**Figure 6.**
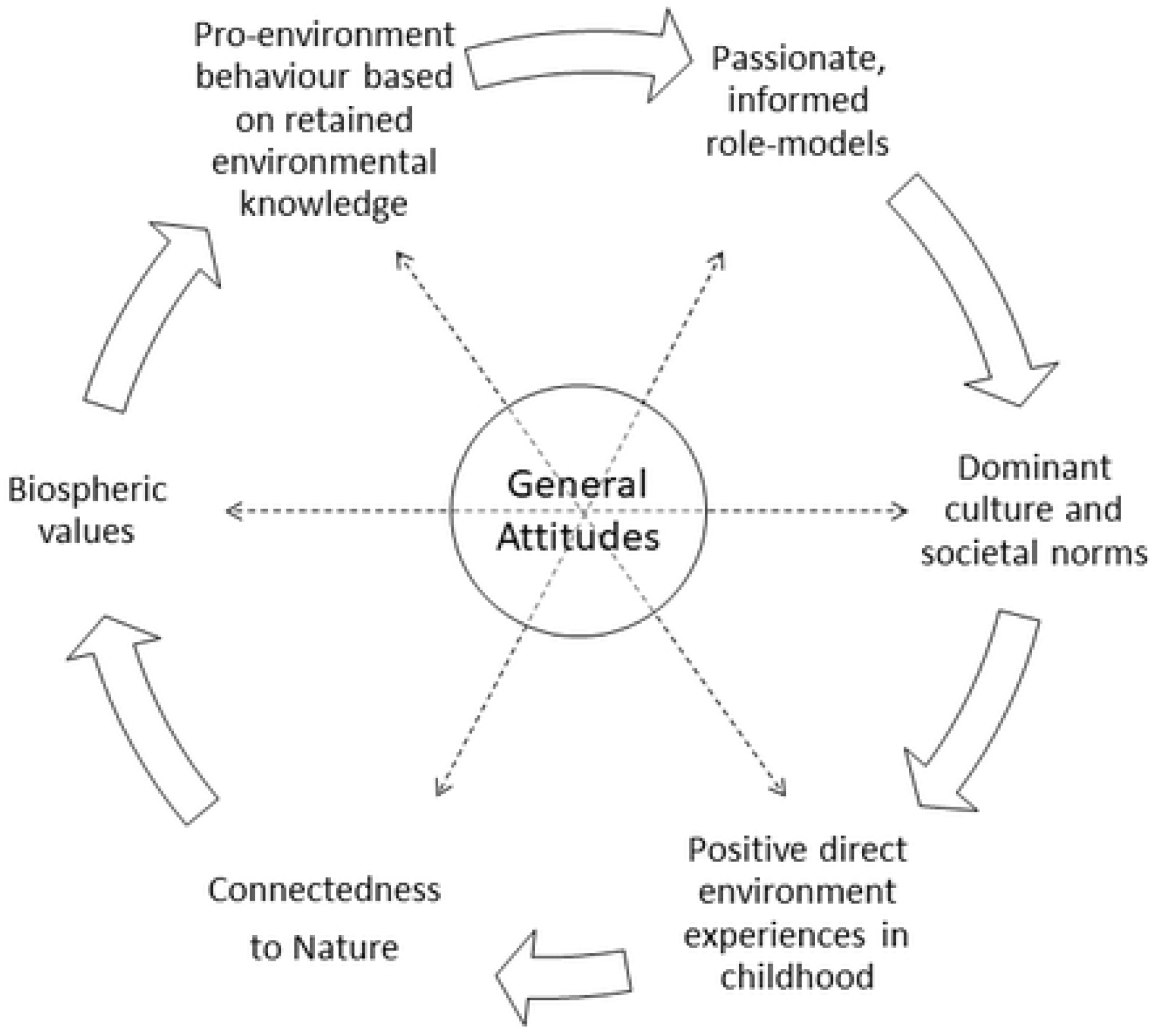
Cycle of knowledge, values and behaviour as the drivers of general attitudes adapted from Klassen et al. (2010).

## Conclusion

This study used a traditionally unpopular group of animals to explore why people like or dislike amphibians and consequently what might motivate them to amphibian stewardship behaviours. It found that individual, charismatic species can be championed amongst groups regardless of affinity towards the class of animals, however positive general attitudes are shaped by a combination of complex social forces, most notably, cultural norms, and regular positive experiences of the species.

